# *In Situ* Graphene-Seq: Spatial Transcriptomics and Chronic Electrophysiological Characterization of Tissue Microenvironments

**DOI:** 10.1101/2025.03.25.645278

**Authors:** Jaeyong Lee, Wenbo Wang, Qiang Li, Zuwan Lin, Ren Liu, Zefang Tang, Junya Aoyama, Richard T. Lee, Xiao Wang, Jia Liu

## Abstract

Biological systems are composed of diverse, interconnected cell types, yet capturing both their functional dynamics and molecular identities at high spatiotemporal resolution remains challenging. While electrophysiological measurements provide real-time insights into cellular activities, they cannot fully describe the molecular architecture and states of the measured cells. Conversely, transcriptomics reveals cell gene expression patterns but does not capture functional states. Bridging these modalities is essential for a holistic understanding of the molecular mechanisms driving functional changes. In this study, we introduce *in situ* graphene-sequencing (graphene-seq), a unique platform that seamlessly integrates chronic electrophysiology with imaging-based, spatially resolved 3D transcriptomics, overcoming longstanding limitations of current multimodal approaches. This system leverages stretchable mesh nanoelectronics for long-term, single-cell-level interfacing and incorporates transparent graphene/PEDOT:PSS electrodes, enabling seamless integration of electrical recordings and optical imaging. By combining electrophysiology with high-throughput, imaging-based *in situ* sequencing, this platform allows comprehensive multimodal, spatially resolved analysis of cell microenvironment within spatially heterogeneous tissues. We validate *in situ* graphene-seq by charting multimodal profiles of human-induced pluripotent stem cell-derived cardiomyocyte and endothelial cell co-cultures, examining how spatial heterogeneity in cell composition influences both electrophysiological activity and gene expression. This scalable, integrated approach offers a powerful tool for studying the complex interplay between cellular function and molecular identity. It also provides insights into how tissue microenvironments shape cell behavior and molecular states, advancing applications in regenerative medicine, stem cell therapy, and disease modeling.

## Introduction

Understanding the relationship between spatially resolved gene expression, cell types, and their microenvironmental niches is fundamental to unraveling the complexities of tissue function^1^. Biological tissues are composed of heterogeneous cell populations that interact dynamically, influencing one another and driving tissue-level functions^1,2,3^. The spatial organization of these cells, their gene expression profiles, and the microenvironmental niches they inhabit—collectively referred to as the cell microenvironment— play critical roles in processes such as development, regeneration, and disease progression^4,5,6,7,8^. Mapping spatially resolved gene expression together with cell types and their niches allows the identification of how microenvironmental differences contribute to molecular and functional heterogeneity within tissues. These findings are important for fields such as neuroscience, cancer biology, and regenerative medicine, and could lay the foundation for targeted therapeutic strategies.

However, linking spatially distinct molecular features to functional outputs such as electrical activity remains technically challenging. Electrical recording methods have uncovered cellular responses and electrophysiological dynamics at the single-cell level^9,10^. For example, high-density electrode arrays with single-cell resolution have been instrumental in probing the coordinated cellular activities underlying sensory, motor, and cognitive functions^11,12,13^. Moreover, long-term monitoring of large ensembles of cells at single-cell resolution has elucidated how cellular activities and connectivity evolve in response to environmental stimuli, development, and aging^14,15^. These high-spatiotemporal-resolution recordings are made possible by engineering devices to seamlessly integrate with tissues^14,15,16^. Despite their utility, electrophysiological recordings alone lack the molecular context necessary to describe the specific types and states of cells underlying these functional changes. Spatial transcriptomics, on the other hand, has been widely employed to characterize cell types and states under spatial context^17^. However, the profiling of the molecular features at fixed samples cannot provide enough insight on the functional behaviors of the cells. Synergistic approaches that combine these modalities have enabled precise cell-type classification and revealed correlations between gene expression and function^18,19,20,21,22,23^. However, existing techniques for multimodal data acquisition are limited in their spatial and temporal resolution. Integrating electrophysiology with transcriptomics at high spatiotemporal resolution can thus provide deeper insights into the molecular mechanisms underlying functional changes.

To address these limitations, we previously introduced *in situ* electro-sequencing, which combines chronic electrophysiology with spatially resolved transcriptomics^24^. Using tissue-like mesh electronics, this platform enables long-term stable, single-cell-level electrical recordings within 3D microtissues. Imaging-based *in situ* sequencing enables extraction of gene expression profiles from electrically recorded cells^25^.

This multimodal approach allows high-throughput mapping of both electrophysiological and transcriptomic states at the single-cell level. However, the opaque metal electrodes in mesh electronics introduce challenges for integrating electrical and optical modalities. Imaging recorded cells on non-transparent electrodes requires additional experimental steps, such as detaching and flipping tissues, which can disrupt the tissue microenvironment. Moreover, as mesh electronics accommodate the dynamic stretching and bending of cellular networks during development, opaque electrodes significantly limit the efficiency of imaging data acquisition within complex 3D structures, such as organoids and spheroids.

Here, we introduce *in situ* graphene-sequencing (graphene-seq), a next-generation *in situ* multimodal profiling platform that seamlessly integrates chronic electrophysiology with high-throughput optical profiling technologies (Fig. 1a). The stretchable mesh structure enables long-term, single-cell-level interfacing across tissues, facilitating chronic electrophysiological analyses (Fig. 1b). By incorporating transparent graphene/PEDOT:PSS electrodes into the mesh, the platform maintains optical clarity, enabling direct imaging of cells on top of the electrodes regardless of their orientation.

**Fig. 1:**
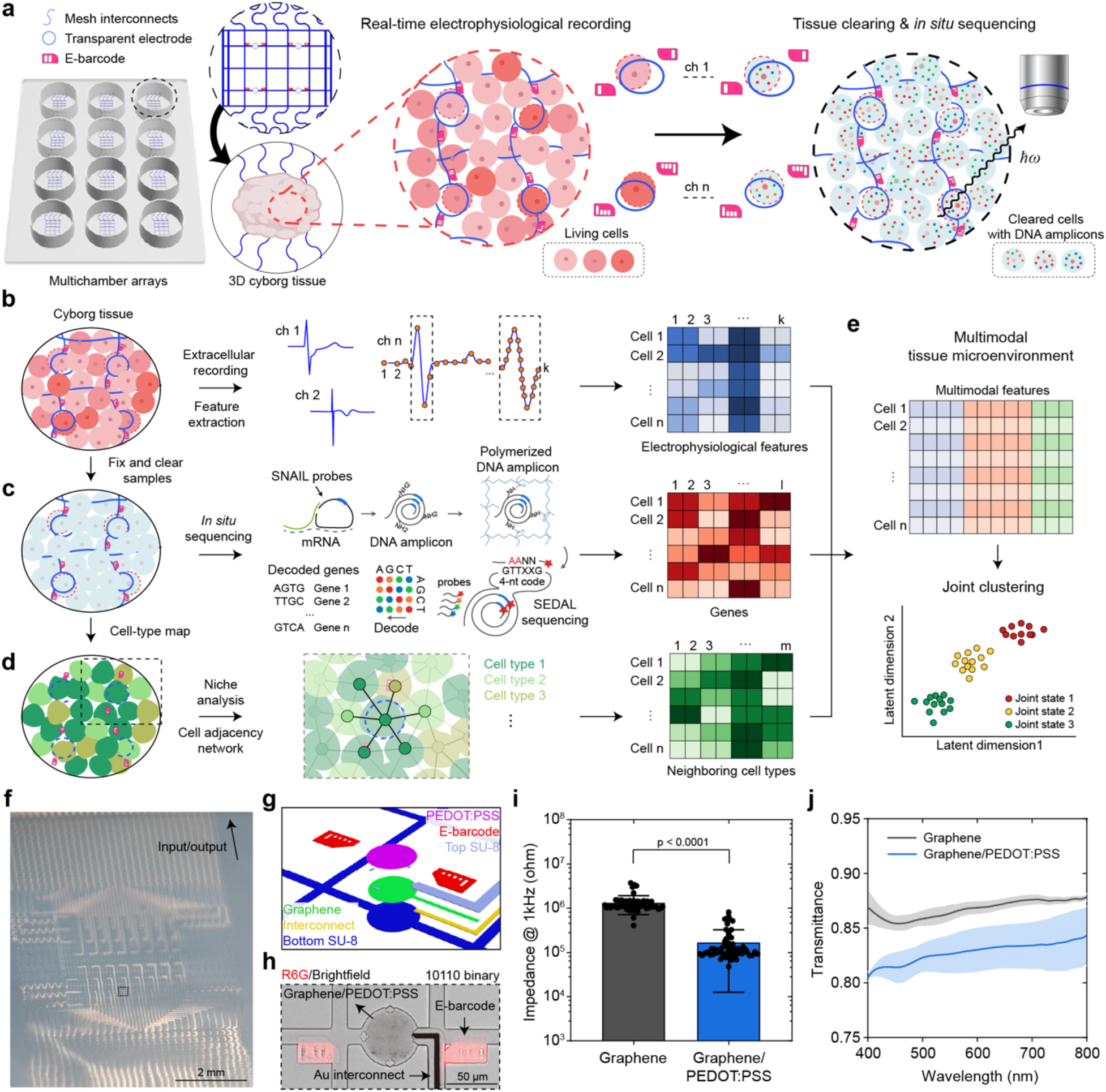
Overview of *in situ* graphene-seq for multimodal tissue analysis. **a**–**e**, Schematics of the *in situ* graphene-seq workflow. **a**, Arrays of stretchable mesh nanoelectronics with transparent graphene/PEDOT:PSS electrodes are integrated with stem cell-derived microtissues for chronic electrophysiological recordings, followed by tissue clearing and *in situ* sequencing. Fluorescent E-barcodes, patterned via photolithography, enable the alignment of electrophysiological recordings with spatial gene expression mapping. **b**, Multichannel extracellular recordings are processed to extract electrophysiological features. **c**, *In situ* sequencing generates spatially resolved single-cell gene expression profiles. **d**, Cell-type maps and niche analyses are constructed based on spatial proximity and cell-type composition. **e**, Joint embedding and clustering of multimodal features integrates electrophysiological, molecular, and cell niche data, providing a comprehensive understanding of tissue microenvironments. **f**, Photograph of a representative mesh nanoelectronics array with transparent electrodes. **g**, Schematic of the electrode structure. **h**, Bright-field (BF) image of a representative graphene/PEDOT:PSS electrode. **i**, Electrochemical impedance of electrodes at 1kHz, measured before and after PEDOT:PSS deposition. Values are presented as mean ± standard deviation (S.D.), p-values were calculated from two-tailed paired t-test, n=61. Each dot represents the impedance of a single electrode within the sample. **j**, Optical transmittance of representative electrodes before and after PEDOT:PSS deposition. Values are presented as mean ± S.D., n=5.

After electrophysiology recording, imaging-based *in situ* sequencing is employed to extract gene expression profiles (Fig. 1c). Reconstructed cell-type maps provide molecularly defined cell types of electrically recorded cells, as well as the local cell composition networks, allowing the study of functional and molecular phenotypes within specific microenvironmental niches (Fig. 1d). Fluorescent E-barcodes, fabricated adjacent to the electrodes, bridge electrophysiological recordings with imaging data, facilitating comprehensive multimodal profiling of tissue microenvironments for further analyses (Fig. 1e). This integrated platform supports the entire workflow—from culturing tissues with devices and recording electrical activities to tissue clearing and high-resolution optical imaging—offering a scalable solution for high-throughput, multimodal profiling on a single platform. By overcoming the optical constraints of previous devices, *in situ* graphene-seq enables, for the first time, seamless and spatially precise correlation of cellular function with gene expression in intact 3D tissues.

## Results

### Design and fabrication of stretchable and transparent mesh nanoelectronics

The stretchable and transparent mesh nanoelectronics consist of a serpentine mesh–structured SU-8 passivation layer and gold (Au) interconnects, as described in previous reports^15,16^. (Fig. 1f and Extended Data Fig. 1a). To incorporate transparent electrodes into the mesh platform, we transferred and patterned 3–5 layers of graphene on top of the interconnects (Methods, Extended Data Fig. 1b,c). The top SU-8 layer passivates the interconnects, exposing only the graphene electrodes for electrophysiological recordings. We fabricated fluorescent E-barcodes next to each electrode for pairwise mapping of electrophysiology and imaging data (Extended Data Fig. 1d–f).

The electrochemical impedance of electrodes is crucial for the signal-to-noise ratio (SNR) in electrophysiological recordings^26^. While graphene microelectrodes offer transparency, they suffer from high electrochemical impedance^27,28,29^. To decrease their impedance while maintaining transparency, we electrochemically deposited PEDOT:PSS on top of the graphene electrodes^30^ (Fig. 1g,h). Deposition was precisely controlled by maintaining a constant current density and adjusting deposition time (Extended Data Fig. 2a,b). Since there is a trade-off between transparency and impedance—both transmittance and electrochemical impedance decrease with increasing deposition time—we optimized the deposition time to balance these parameters (Methods, Extended Data Fig. 2c–f). This optimization reduced the electrochemical impedance at 1 kHz from 1,319 ± 600 kΩ to 168.7 ± 156.1 kΩ after depositing PEDOT:PSS (mean ± standard deviation (S.D.), Fig. 1i), while maintaining transmittance above 80% across the visual spectrum, making it compatible with most bioimaging techniques (Fig. 1j). Furthermore, the graphene/PEDOT:PSS electrodes demonstrate stable electrochemical performance across different samples (Extended Data Fig. 2g) and maintained their performance for over 60 days of incubation in 37 ℃ 1X PBS solution (Extended Data Fig. 2h,i).

The fabrication process is highly scalable, with all steps compatible with conventional 2D microfabrication techniques. This adaptability allows straightforward modifications of the design to accommodate specific experimental setups, such as multi-well cell plates, which are commonly used for tissue culture (Extended Data Fig. 3). Additionally, recent advancements in graphene manufacturing and transfer techniques enable the mass production of graphene-integrated devices^31,32,33^, enhancing the potential for broad and customizable applications.

### Electrical and optical performance within 3D organoids

We next integrated stretchable and transparent mesh electronics with 3D cardiac organoids using established methods (Methods, Extended Data Fig. 4a–c)^16^. Initially, the devices were integrated with 2D sheets of human-induced pluripotent stem cell (hiPSC)-derived cardiomyocytes (CMs) cultured on a Matrigel layer. During differentiation and self-organization, the transition from 2D to 3D in the formation of cardiac organoids folded the devices into 3D structures, ultimately embedding them within the organoids.

To confirm the integration and transparency of graphene/PEDOT:PSS electrodes within organoids, we performed tissue fixation, clearing, and immunostaining, followed by confocal microscope imaging (Methods, Fig. 2a,b). The 3D reconstructed images show the mesh electronics forming interwoven structures within the cellular networks (Fig. 2a), while bright-field (BF) images show the spatial distribution of recording electrodes across the organoids (Fig. 2b). Unlike the metal interconnect layer, which blocks optical signals, the graphene/PEDOT:PSS electrodes allow imaging of fluorescence markers directly above the electrodes (Fig. 2c,d).

**Fig. 2:**
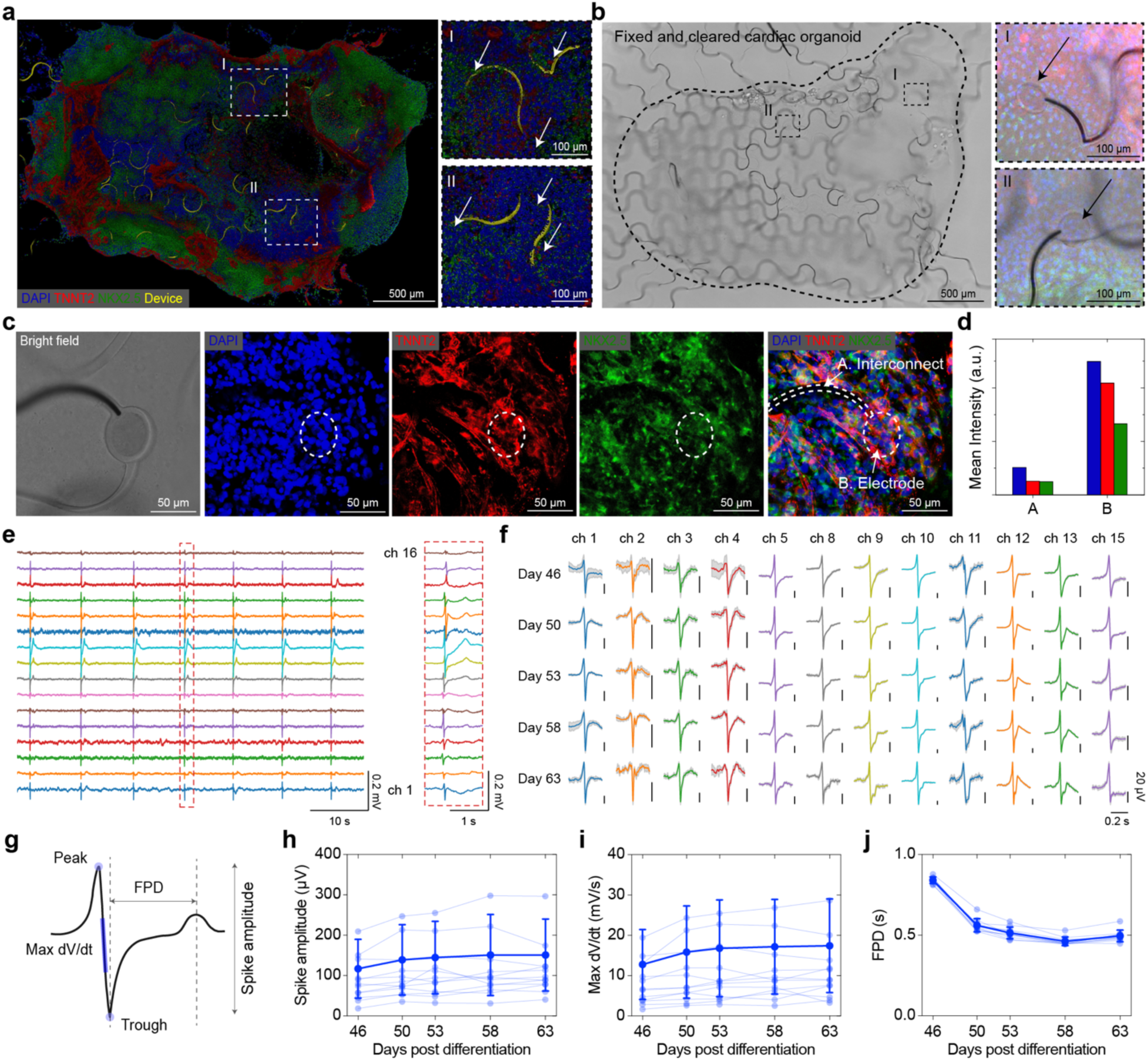
3D cardiac organoid integration and long-term electrophysiological recordings. **a**–**b**, 3D reconstructed images of cardiac organoids integrated with mesh nanoelectronics. **a**, Confocal fluorescence reconstruction. Right panels provide zoomed-in views of the dashed-box regions in the left panel. White arrows highlight meshes embedded within the organoid. **b**, BF images. Right panels provide zoomed-in views of dashed-box regions in the left panel. Black arrows indicate transparent graphene/PEDOT:PSS electrodes. **c**, BF and confocal fluorescence images of immunohistochemically stained cardiomyocytes surrounding a representative graphene/PEDOT:PSS electrode. Regions A and B indicate areas over metal interconnect and graphene/PEDOT:PSS electrode, respectively. **d**, Quantification of mean pixel intensity for three fluorophores over the metal interconnect region (A) and the graphene/PEDOT:PSS electrode region (B) shown in (**c**). **e**, Representative voltage traces recorded from 16 electrodes. The right panel shows zoomed-in spike waveforms. **f**, Spike waveforms (mean ± S.D.) extracted from the same electrodes across five recording sessions. **g**, Schematic of waveform features extracted for statistical analysis: spike amplitude, maximum dV/dt, and field potential duration (FPD). **h**–**j**, Temporal tracking of waveform features across five recording sessions: spike amplitude (**h**), maximum dV/dt (**i**), and FPD (**j**). Values are presented as mean ± S.D. Sample sizes: (**h**) n=12; (**i**) n=12; (**j**) n=10. Each dot represents a feature extracted from the mean waveform of a single electrode.

Previous approaches relied on opaque metal electrodes required additional procedures, such as detaching the tissue/electronics hybrid, flipping the sample, and transferring it to another plate for imaging electrically recorded cells (Extended Data Fig. 4d–f). During these processes, the cell microenvironment cannot be fully preserved. Moreover, the direction of electrodes affected image acquisition of recorded cells. By incorporating transparent graphene/PEDOT:PSS electrodes, we eliminated these additional steps and enabled both electrophysiological recordings and imaging on a single platform, while preserving the original spatial localization of the tissue (Extended Data Fig. 4g–i). The use of imaging-quality glass wafers further enhanced imaging clarity, improving overall efficiency and streamlining experimental workflows. These improvements allow unobstructed optical interrogation of electrically recorded cells regardless of electrode directions, facilitating high-throughput multimodal analyses within 3D tissues.

Next, we evaluated the recording stability of the distributed graphene/PEDOT:PSS electrodes in hiPSC-derived cardiac organoids. Voltage traces from 16 representative channels, recorded 22 days after device integration, are shown in Fig. 2e. Recordings were performed across five sessions using the same organoid, during which mean spike waveforms from effective channels were extracted (Fig. 2f). The graphene/PEDOT:PSS electrodes demonstrated stable extracellular field potential recordings throughout the organoid over time. Furthermore, the stretchable and transparent mesh electronics facilitated the tracking of waveform features, including spike amplitude, absolute maximum dV/dt (maximum dV/dt), and field potential duration (FPD), across 3D organoids (Fig. 2g). No significant changes were observed in spike amplitudes or maximum dV/dt over time (Fig. 2h,i), while a reduction in FPD was noted between days 46 and 50 post-differentiation (Fig. 2j). These results highlight the long-term recording stability of the stretchable and transparent mesh electronics, enabling detailed monitoring of electrophysiological dynamics in 3D tissue models.

### Multimodal charting of electrophysiology, transcriptomics, and cell niches

*In situ* graphene-seq streamlines the multimodal data acquisition process by seamlessly integrating electrophysiological recordings with imaging-based *in situ* sequencing. To demonstrate this, we first integrated hiPSC-derived cardiac microtissues with stretchable and transparent mesh electronics to record electrophysiological signals. Extracellular spike waveforms were collected, aligned to each channel, and processed to extract electrophysiological features (Methods).

Immediately after recordings, the tissue/electronics hybrid was fixed, followed by *in situ* complementary DNA (cDNA) library preparation and sequencing to capture gene expression profiles within a spatial context (Methods). Briefly, the recorded tissue was fixed with 4% paraformaldehyde (PFA) and subjected to *in situ* conversion of messenger RNA (mRNA) from targeted genes into a cDNA library. The tissue was then embedded in hydrogel, cleared, and processed using previously developed sequencing-by-ligation scheme to decode gene identities through multiple rounds of imaging^24,25^. Fluorescent E-barcodes positioned near each electrode were imaged alongside DNA amplicons, allowing precise alignment of the *in situ* sequencing data with electrical recordings.

We then applied ClusterMap for 3D cell segmentation and amplicon read assignment to individual cells^34^ (Methods, Extended Data Fig. 5). After segmentation and data preprocessing, we performed *Leiden* clustering^35^ on Uniform Manifold Approximation and Projection (UMAP) embeddings^36^ to identify major cell types. Electrically recorded cells were identified by aligning segmented cells with electrodes imaged in reflection mode. Electrode segmentation was achieved using the Segment Anything Model (SAM)^37^, a pretrained visual foundation model for accurate generation of electrode masks that isolated electrodes and reconstructed them as 3D point clouds. Cells recorded by the electrodes were identified by calculating their proximity to the convex hull center derived from the reconstructed electrode point clouds. Representative multimodal data—including electrophysiological recordings, gene expression profiles, adjacent cell network analysis, and the combined multimodal features of hiPSC-derived cardiac microtissues at day 30 post-differentiation—are shown in Extended Data Fig. 6.

### Capturing functional and molecular heterogeneity within spatially heterogeneous tissue

*In situ* graphene-seq can simultaneously capture electrophysiological signals and molecular profiles from complex 3D tissues, making it well-suited for investigating niche-specific effect within spatially heterogeneous environments. To demonstrate this, we have explicitly built the spatially heterogeneous tissue model by co-culturing hiPSC-derived CMs (hiPSC-CMs) and hiPSC-derived endothelial cells (hiPSC-ECs) in varying ratios within a single device. Co-culturing CMs with ECs is known to enhance the electrical maturation of CMs, improving metrics such as spike amplitude, maximum dV/dt, and spike duration^23^. However, how local variations in cell composition within the tissue impact these functional properties remains unclear. To address this, we created a heterogeneous microtissue model by co-culturing hiPSC-CMs and hiPSC-ECs at a 3:1 ratio on one side of the mesh electronics, while seeding hiPSC-CMs only on the opposite side (Methods, Fig. 3a). Immunohistostaining for cardiac troponin T (TNNT2) and cluster of differentiation 31 (CD31) revealed distinct cellular distributions, with higher densities of CD31-positive hiPSC-ECs in the CM/EC co-culture region (Fig. 3b).

**Fig. 3:**
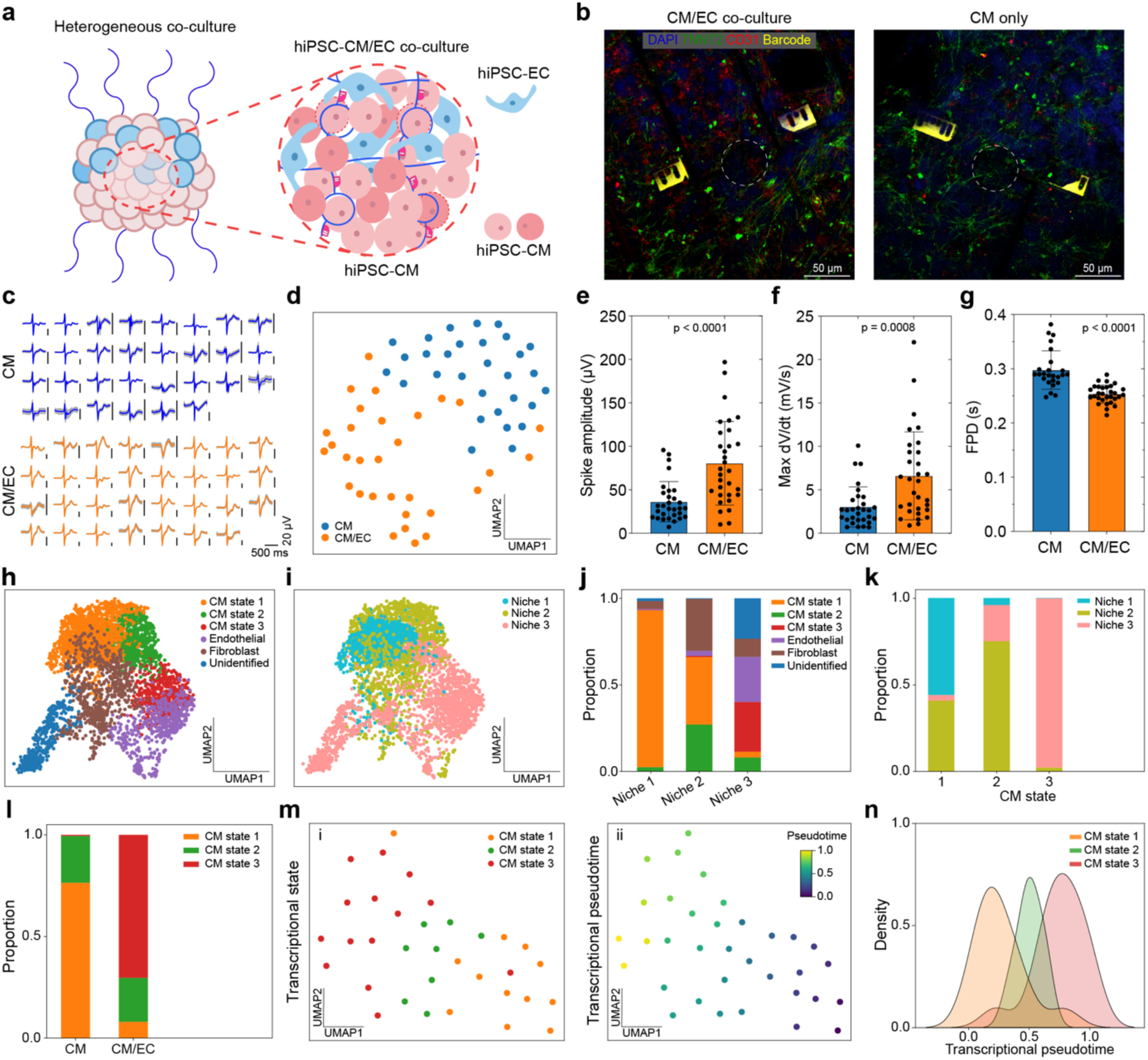
Spatial heterogeneity in hiPSC-CM/hiPSC-EC co-cultured microtissues. **a**, Schematic illustration of spatially heterogeneous cell culture, with hiPSC-CMs co-cultured with hiPSC-ECs (CM/EC) on one side of the device and hiPSC-CMs cultured alone (CM) on the other side. **b**, Confocal fluorescence images showing immunohistochemically stained cells near electrodes in the CM/EC co-culture region and the CM only region. **c**, Extracted spike waveforms (mean ± S.D.) from electrodes in the CM/EC and CM only regions of the tissue. **d**, Uniform Manifold Approximation and Projection (UMAP) embedding of spike waveforms, color-coded by electrode location. **e**–**g**, Statistical comparison of waveform features between the CM/EC and CM only regions, including spike amplitude (**e**), maximum dV/dt (**f**), and FPD (**g**). Values are presented as mean ± S.D., p-values were calculated from two-tailed unpaired t-test. Sample sizes: (**e**) CM (n=30), CM/EC (n=31); (**f**) CM (n=30), CM/EC (n=31); (**g**): CM (n=25), CM/EC (n=30). Each dot represents a feature extracted from the mean waveform of a single electrode. **h–i**, Classification of hiPSC-CM states and cell types (**h**) and molecular niches (**i**) based on transcriptional profile. **j**, Bar plots showing the average proportions of cell types and states within each molecular niche. **k**, Distribution of molecular niches associated with different hiPSC-CM states. **l**, Distribution of hiPSC-CM states in the CM only and CM/EC regions. **m**, UMAP embedding of gene expression profiles of electrically recorded hiPSC-CMs, color-coded by cell states (i) and transcriptional pseudotime (ii). **n**, Density plot showing the distribution of transcriptional pseudotime values across different hiPSC-CM states.

To compare electrophysiological characteristics across the regions, we recorded electrophysiological signals, extracted waveforms, and aligned them with electrode positions. Channels 1–32 recorded signals from the CM/EC co-culture region, while channels 33–64 captured data from the CM-only region (Fig. 3c). Projecting the mean waveforms from each channel into a 2D UMAP embedding revealed distinct clusters corresponding to the CM-only and CM/EC co-culture regions (Fig. 3d). Waveform feature analysis indicated enhanced electrical maturation of hiPSC-CMs in the CM/EC co-culture region, evidenced by higher spike amplitudes, greater maximum dV/dt, and shorter FPD compared to the CM-only region (Fig. 3e–g).

*In situ* sequencing revealed diverse CM states and additional cell types, such as endothelial cells and fibroblasts (Fig. 3h). Cell niches were characterized by analyzing the local cellular composition surrounding each cell, which was organized into a cell-by-composition matrix for dimensionality reduction and clustering (Methods). This analysis identified three primary cell niches, each defined by distinct cell-type proportions (Fig. 3i,j). Notably, specific CM states were associated with different niches: CM state 3 (CM_s3) was predominantly enriched in niche 3, while CM state 1 (CM_s1) was primarily localized in niche 1 (Fig. 3k). Additionally, CM-only and CM/EC co-culture regions displayed distinct proportions of cell states, with CM_s1 being enriched in CM-only regions and CM_s3 in CM/EC co-culture regions (Fig. 3l). CM state 2 (CM_s2) was present in both CM-only and CM/EC co-culture regions.

Gene expression analysis highlighted distinct molecular characteristics among CM states (Extended Data Fig. 7). CM_s1 exhibited higher expression of genes such as *PDK1* and *HAND2*, which are associated with early developmental stages^38,39^. In contrast, CM_s2 and CM_s3 showed upregulation of ion channel genes critical for excitation-contraction coupling and functional maturation. Notably, potassium channel genes, including *KCNE2*, *KCNK3*, and *KCNA4*, were highly expressed, along with *PIEZO1*, a mechanosensitive ion channel involved in responses to mechanical stress^40^. CM_s3 displayed robust expression of structural and adhesion-related genes, such as *VCL*, *ITGB5*, and *CDH2*, as well as gap junction-related genes, including *GJA1* (Connexin 43) and *GJA5* (Connexin 40), suggesting advanced functional maturity.

To further explore molecular maturation, we performed pseudotime trajectory analysis, which estimates continuous cell states based on transcriptomic or electrophysiological data (Methods, Fig. 3m)^41,42,43^. The results showed a clear progression along the pseudotime trajectory, with molecular pseudotime sequentially increasing from CM_s1 to CM_s2 and finally to CM_s3 (Fig. 3n). The observed alignment between electrical and molecular heterogeneity in hiPSC-CMs across different tissue regions highlights the capability of *in situ* graphene-seq to capture local microenvironmental influences on both the functional and molecular characteristics of cells within the tissues.

### Integrative analysis of functional and molecular characteristics in spatially resolved cell niches

Spatially resolved *in situ* RNA sequencing enables precise cell-type classification and niche analysis, facilitating the identification and classification of tissue regions^8,44^. *In situ* graphene-seq can extend this capability by integrating functional characterization of cells within molecularly defined regions (Fig. 4a), allowing the study of functional distinctions driven by cellular microenvironments.

**Fig. 4:**
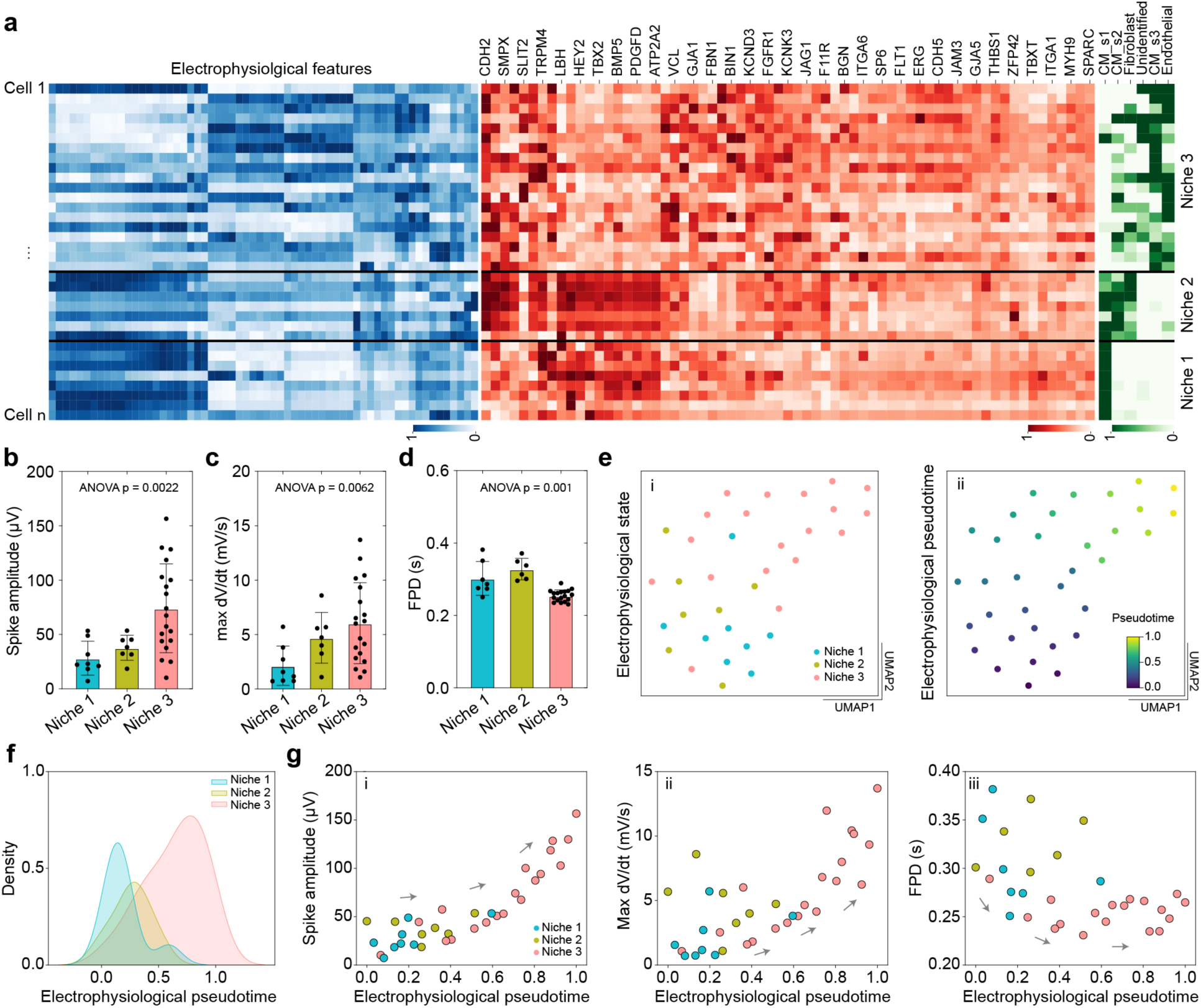
Electrophysiological characterization of molecularly defined cell niches. **a**, Concatenated heatmaps displaying electrophysiological features, selected gene expressions, and local cell compositions of identified cells, grouped by molecularly defined cell niches. **b**–**d**, Comparative analysis of waveform features in CMs across the three cell niches: spike amplitude (**b**), maximum dV/dt (**c**), and FPD (**d**). Data are presented as mean ± S.D., p-values were calculated from ordinary one-way ANOVA. Sample sizes: (**b**) Niche 1 (n=8), Niche 2 (n=7), Niche 3 (n=19); (**c**) Niche 1 (n=8), Niche 2 (n=7), Niche 3 (n=19); (**d**) Niche 1 (n=7), Niche 2 (n=6), Niche 3 (n=18). Each dot represents a feature extracted from the mean waveform of a single electrode. **e**, UMAP embedding of electrophysiological features from electrically recorded CMs, color-coded by cell niche (i) and electrophysiological pseudotime (ii). **f**, Density plot showing the distribution of electrophysiological pseudotime values across the three cell niches. **g**, Scatter plots showing correlations between electrophysiological pseudotime and waveform features: spike amplitude (i), maximum dV/dt (ii), and FPD (iii). Each dot represents spike features from a single electrode and is color-coded by molecularly defined cell niche. Increased pseudotime values are associated with higher spike amplitude, greater maximum dV/dt, and shorter FPD of hiPSC-CMs, indicating a progression toward more electrically matured states.

We first analyzed waveform features of electrically recorded cells within different molecularly defined niches (Fig. 4b–d). Spike amplitude, maximum dV/dt, and FPD exhibited distinct characteristics across Niche 1, Niche 2, and Niche 3, reflecting functional differences across these regions. Notably, spike amplitude and maximum dV/dt progressively increased from Niche 1 to Niche 2 and Niche 3. To further examine these trends, we performed pseudotime trajectory analysis on embeddings of electrophysiological features extracted from the waveforms (Methods, Fig. 4e). A density plot showed that cells in Niche 3 clustered at higher electrophysiological pseudotime trajectory values, followed by cells in Niche 2 and Niche 1 (Fig. 4f). Scatterplots correlating electrophysiological pseudotime with waveform features demonstrated increasing spike amplitude, greater maximum dV/dt, and shorter FPD with advancing electrophysiological pseudotime (Fig. 4g). These findings suggest that CMs in Niche 3 represent a more electrically matured state, followed by those in Niche 2 and Niche 1. Additionally, the results closely align with molecular pseudotime trajectories, where Niche 1 is predominantly associated with CM_s1, Niche 2 with CM_s2, and Niche 3 with CM_s3. This highlights the capability of *in situ* graphene-seq to capture functional distinctions driven by molecularly defined niches.

*In situ* graphene-seq integrates spatial mapping of gene expression, cell-type distributions, and electrophysiological recordings, providing a comprehensive framework for understanding tissue microenvironments. Representative electrodes positioned on cell-type maps of distinct niches, along with the corresponding waveforms recorded from these electrodes, highlight the molecular and functional heterogeneity across tissue regions (Fig. 5a–c). By leveraging these multimodal datasets, our platform suggests a more comprehensive approach of tissue organization, expanding current molecular-based criteria into an integrative multimodal framework. We demonstrated the joint definition of cell niches by combining electrophysiology, gene expression, and local cell composition data (Fig. 5d,e). The joint embedding of concatenated multimodal features is shown in Fig. 5d and 5e, with color-coding indicating the identified joint niches (Fig. 5d) and tissue regions—CM only or CM/EC co-culture (Fig. 5e).

**Fig. 5:**
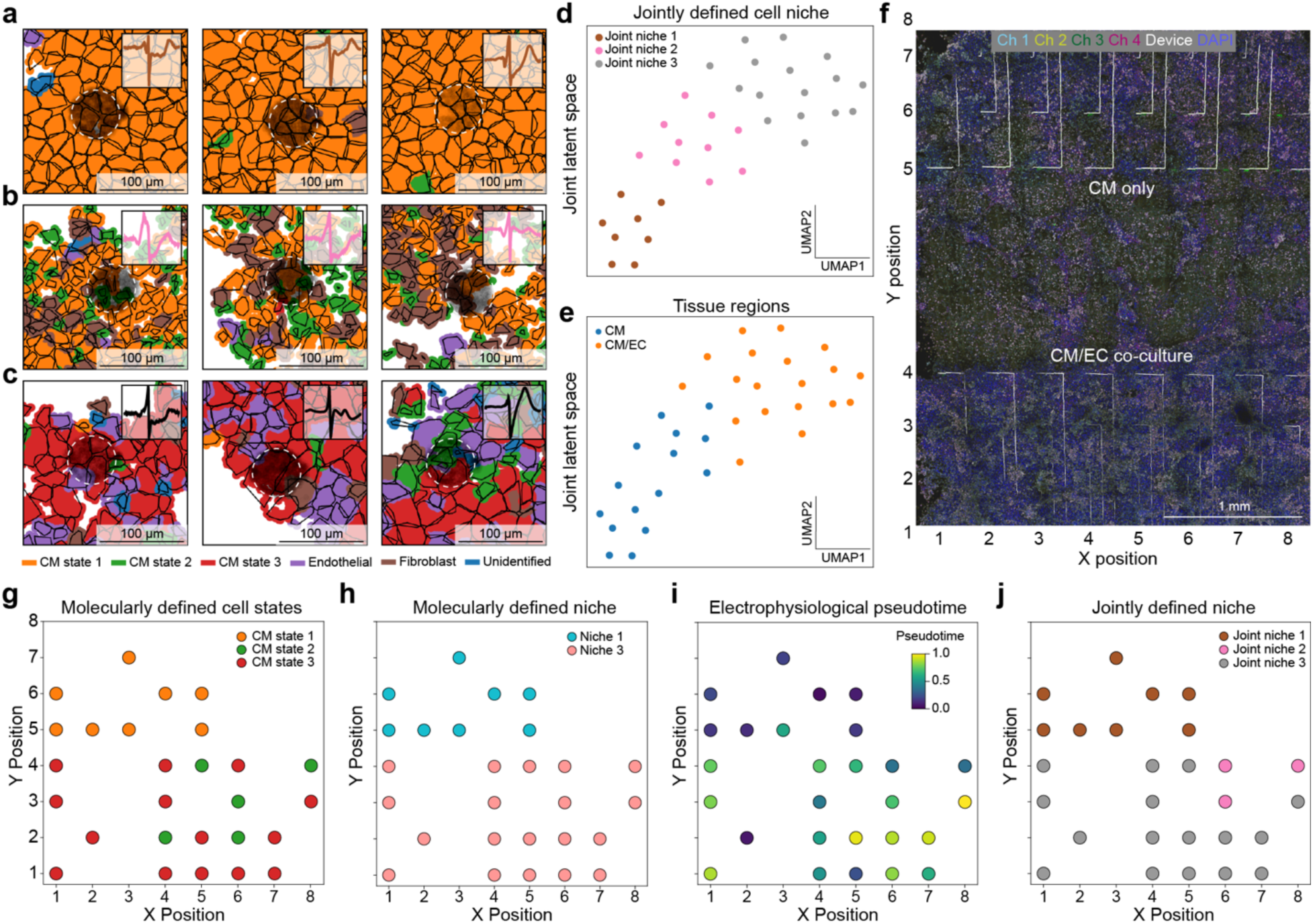
Multimodal cell niches and tissue annotation. **a**–**c**, Representative cell-type maps of electrically recorded CMs in molecularly defined niche 1 (**a**), niche 2 (**b**), and niche 3 (**c**) with their recorded waveforms overlapped. **d**–**e**, Joint embedding of electrophysiological features, gene expression, and local cell composition into 2D UMAP spaces, color-coded by jointly defined cell niches (**d**) and tissue regions (CM only or CM/EC co-cultured regions) (**e**). **f**, Representative *in situ* sequencing image of a heterogeneous microtissue containing CM only and CM/EC regions, with relative X and Y positions of electrodes. **g**–**j**, Spatial mapping of electrode locations across the tissue in (f). Each dot represents an electrode, color-coded by the transcriptional state of the electrically recorded hiPSC-CMs (**g**), molecularly defined cell niche (**h**), electrophysiological pseudotime (**i**), and jointly defined cell niche (**j**).

Since our platform allows precise localization of electrodes within the tissue model (Fig. 5f), spatial information can be directly mapped back onto the original tissue architecture, providing an integrated perspective on the relationship between spatial organization and functional properties. By identifying the relative positions of electrodes within the sample, we defined regional profiles in heterogeneously cultured tissues, facilitating a detailed investigation of molecularly defined cell niches, cell states, and electrophysiological traits, as well as the precise classification of the multimodal cell microenvironments (Fig. 5g–j).

## Conclusion

In this study, we present *in situ* graphene-seq, a platform that employs stretchable and transparent mesh nanoelectronics to seamlessly integrate with 3D organoids or microtissues. Fabricated directly on imaging glass wafers, the mesh electronics incorporate transparent electrodes that facilitate high-resolution imaging directly over the electrodes, regardless of their orientation. This design allows for simultaneous electrical recordings and imaging-based spatial transcriptomics within 3D-integrated tissues while preserving their original architecture

Using *in situ* graphene-seq, we captured rich multimodal datasets from spatially heterogeneous microtissues, enabling the identification and analysis of specific niches or microenvironments. We identified distinct electrophysiological properties and molecular signatures of cells based on their microenvironments. Multimodal cell-niche analysis, guided by spatial proximity and cell-type composition, further revealed how electrophysiological features and cell states are influenced by neighboring cell compositions. Additionally, we showed how this integrative multimodal framework provides spatial, molecular, and functional insights into tissue organization. Cross-modal prediction models, such as those using long-term electrical recordings to infer molecular cell states or niches, may further illuminate dynamic interactions within heterogeneous tissues.

There are several exciting directions to extend the capabilities of *in situ* graphene-seq. One opportunity is to incorporate real-time or longitudinal monitoring of multimodal phenotypes and integrate those data with *in situ* molecular analyses. For example, a recent study has shown that graphene field-effect transistors can function as multifunctional biosensors, simultaneously measuring electrophysiological and mechanophysiological signals^45^, providing the potential for incorporating the mechanical contraction forces as a unique modality to jointly define the cell state. In addition, advances in spatial transcriptomics promise to extend the capability of *in situ* sequencing to the realm of multi-omics, including spatial transcriptomics with proteins imaging^46^, transcriptome-translatome co-profiling, and sequencing of intact organoids or tissue blocks^47^. Applying *in situ* graphene-seq *in vivo* could further enable the investigation of multimodal characteristics in brain and cardiac systems at high spatiotemporal resolution, allowing the comprehensive multimodal profiling of the systems. Furthermore, the integration of large-scale biological datasets, such as single-cell gene expression data, with artificial intelligence (AI) models is transforming our ability to interpret biological and biomedical contexts from the data^48,49,50^. With its scalable design, our platform is well-suited for high-throughput data acquisition, facilitating the collection of large multimodal datasets. We envision that these advancements position *in situ* graphene-seq as a powerful tool for multimodal tissue analysis, offering comprehensive insights into the complex biological systems.

## Methods

### Device fabrication

Fabrication steps of stretchable and transparent mesh nanoelectronics are as follows: (1) A 3-inch glass wafer (D263 Glass wafer, UniversityWafer) was cleaned using a piranha solution (3:1 mixture of sulfuric acid and 30% hydrogen peroxide), followed by rinsing with acetone, isopropyl alcohol (IPA), and deionized (DI) water. (2) Nickel (Ni) sacrificial layer patterning: Hexamethyldisilazane (HMDS, MicroChem) and LOR 3A (MicroChem) were spin-coated at 3000 rpm and baked at 180 ℃ for 5 minutes. Photoresist S1805 (MicroChem) was spin-coated at 3000 rpm, baked at 115 ℃ for 1 minute, exposed to 405 nm ultraviolet (UV) light at 40 mJ/cm², and developed using CD-26 developer (Microposit). The sample was descummed using O_2_ plasma (12 sccm flow, 50 W for 1 minute, Samco PC-300). (3) A 100 nm-thick Ni layer was deposited using a Sharon thermal evaporator, followed by a standard lift-off process in remover PG (MicroChem). (4) Bottom SU-8 patterning: SU-8 2002 (MicroChem) was spin-coated at 4000 rpm, pre-baked at 65 ℃ for 2 minutes and 95 ℃ for 4 minutes, and exposed to 365 nm UV light at 200 mJ/cm². The sample was post-baked at 65 ℃ and 95 ℃ for 2 minutes each, developed using SU-8 developer (MicroChem), and then hard-baked at 180 ℃ for 1 hour. (5) Metal interconnect patterning: A lift-off pattern was created as described in step (2). Gold (Au) with chromium (Cr) adhesive layers (Cr/Au/Cr, 5/70/5 nm-thick) was deposited using an e-beam evaporator (Denton) and lifted off in remover PG (MicroChem). (6) Graphene wet transfer: 950 PMMA A4 (Kayaku Advanced Materials) was spin-coated at 3000 rpm on graphene on copper foil (ACS Material) and baked at 150 ℃ for 3 minutes. The backside graphene was optionally etched using O_2_ plasma (12 sccm flow, 50 W for 1 minute). Copper was etched using a copper etchant (APS-100, Transene) for 2 hours, followed by rinsing with DI water for three times, 30 minutes each. The freestanding graphene/PMMA film was transferred to the wafer and baked at 80 ℃ overnight. The PMMA layer was then removed using acetone. (7) Graphene patterning: HMDS and Photoresist S1813 (MicroChem) were spin-coated at 3000 rpm, baked at 115 ℃ for 1.5 minutes, exposed to 405 nm UV light at 150 mJ/cm², and developed using CD-26 developer. Graphene was etched using O_2_ plasma (12 sccm flow, 150 W for 3 minutes). (8) Top SU-8 passivation: Conditions described in step (4) were used. (9) E-Barcode patterning: A 0.004 wt% mixture of Rhodamine 6G (Sigma-Aldrich) and SU-8 2000.5 (MicroChem) was spin-coated at 2000 rpm. The sample was pre-baked at 65 ℃ for 1 minute and 95 ℃ for 3 minutes, exposed to 365 nm UV light at 60 mJ/cm², post-baked at 65 ℃ for 1 minute and 95 ℃ for 2 minutes, developed in SU-8 developer, and hard-baked at 180 ℃ for 1 hour. (10) Flexible flat cables (Molex) were connected to the input/output pads using a flip-chip bonder (Finetech). (11) A customized chamber was glued onto the wafer, and electrical components outside the chamber were encapsulated with a low-toxicity silicone adhesive (Kwik-Sil, World Precision Instrument). (12) The Ni sacrificial layer was released by adding ferric chloride (MG Chemicals) into the chamber, followed by etching for 2–3 hours at 40 ℃ and rinsing with DI water several times. (13) The solution inside the chamber was replaced with EDOT:PSS (0.01 M and 0.1 M in DI water), and PEDOT:PSS was electroplated by applying a DC current (0.4 mA/cm²) for 4 seconds using SP-150 potentiostat (Bio-Logic).

### Electrical and optical characterization

The Electrochemical Impedance Spectra (EIS) of graphene and graphene/PEDOT:PSS electrodes were measured using SP-150 potentiostat. The devices were immersed in 1X PBS with a silver/silver chloride (Ag/AgCl) reference electrode and a platinum counter electrode. A sinusoidal voltage with a 100 mV peak-to-peak amplitude was applied, with frequencies ranging from 1 MHz to 1 Hz. The impedance at 1 kHz of the electrodes was characterized using the integrated functions of the Blackrock CerePlex and Intan RHD Recording systems. Optical transmittance of the graphene and graphene/PEDOT:PSS electrodes was measured using CytoViva hyperspectral microscopy. Data and statistical analyses were performed using GraphPad Prism.

### Cell culture and differentiation

#### Stem cell culture and cardiomyocytes differentiation

Human induced pluripotent stem cells (hiPSCs) from the IMR-90 line were obtained from the WiCell Research Institute (Madison, WI, USA), with mycoplasma testing performed by the vendor. The hiPSCs were maintained and cultured on Matrigel-coated 6-well plates in Essential 8 medium (Gibco). Culture medium was replaced daily to maintain cell pluripotency. Cells were passaged every 4 days using 5 mM EDTA (Invitrogen). Cardiomyocyte differentiation was performed following a previously established protocol^16,24^. Briefly, hiPSCs at approximately 80% confluency were treated with RPMI 1640 (Gibco) supplemented with 2% B27 minus insulin (Gibco) and 12 µM CHIR99021 (BioVision) for 24 hours. Subsequently, cells were incubated in RPMI 1640 with 2% B27 minus insulin for 48 hours, followed by 48 hours of treatment with RPMI 1640 containing 2% B27 minus insulin and 5 µM IWR1(Cayman Chemical). Cells were then cultured in RPMI 1640 with 2% B27 minus insulin for 48 hours before transitioning to RPMI 1640 supplemented with 2% B27 for extended culture. Quality control was performed by observing spontaneous beating of the tissue, typically evident 2-3 days after changing the medium to RPMI 1640 with 2% B27. Culture medium was replaced every 48 hours. hiPSC-derived endothelial cells were cultured in Endothelial Cell Growth Medium-2 (Lonza). The media was changed every 4 days.

#### Integration of hiPSC-derived cardiomyocytes and endothelial cells with devices

For all device integrations, 10 µM ROCK inhibitor (Tocris Bioscience) was added to the cell culture medium for the first 24 hours post-integration and omitted in subsequent cultures. Prior to cell integration, devices were washed twice with 1X PBS. The devices were then incubated in a bind-silane solution (10% v/v acetic acid, 1% v/v bind silane in ethanol) for 3 hours, followed by three washes with 90% ethanol. Subsequently, devices were incubated overnight in 0.01% w/v Poly-D-lysine hydrobromide solution (Sigma-Aldrich), washed three times with DI water, and coated with Matrigel solution (100 µg/mL, Corning) at 37 °C for 1 hour. The devices were then chilled, and 50 µL of 10 mg/mL Matrigel solution was added from the edge. The device/Matrigel hybrid was incubated at 37 °C for 30 minutes.

hiPSC-derived cardiomyocytes (hiPSC-CMs) were dissociated into single cell suspensions by incubation in 0.05% Trypsin-EDTA solution (Gibco) for 5 minutes. Cell viability was assessed using trypan blue staining (Gibco). Approximately 3–4 million of hiPSC-CMs were resuspended in RPMI 1640 medium supplemented with 2% B27 and seeded onto the device/Matrigel hybrid.

For CM/EC heterogeneous co-culture, hiPSC-derived endothelial cells (hiPSC-ECs) were dissociated into single-cell suspensions using TrypLE Express (Gibco) at 37 °C for 3 minutes. hiPSC-CMs and hiPSC-ECs were mixed in a 3:1 ratio and resuspended in StemPro-34 SFM (Gibco) supplemented with 1X StemPro-34 Nutrient Supplement (Gibco), 1X GlutaMAX-I (Gibco), and 50 ng/mL human VEGF165 (GenScript). Two million cells were seeded to cover half of the device chamber. The chambers were tilted to prevent cell diffusion. After 24 hours post-seeding, the remaining half of the device was seeded with 2 million hiPSC-CMs. All heterogeneous cultures were maintained in StemPro-34 SFM supplemented with 1X StemPro-34 Nutrient Supplement, 1X GlutaMAX-I, and 50 ng/mL human VEGF165. Culture medium was changed every 48 hours.

### Immunohistostaining

Immunohistostaining was performed using adapted methods from the previous report^16^. Tissues were fixed with 4% PFA solution (Electron Microscopy Sciences) at 4 ℃ overnight. The samples were then incubated in a hydrogel solution (0.1% VA-044 (Fujifilm Wako) and 4% acrylamide (Bio-Rad) in 1X PBS) at 4 ℃ for 24 hours, followed by hydrogel polymerization at 37 ℃ for 3 hours. The polymerized samples were incubated in Electrophoretic Tissue Clearing solution (Logos Biosystems) at 37 ℃ for 48 hours and subsequently rinsed several times with PBST (1X PBS + 0.1% Tween-20 (Millipore)). The samples were then incubated with primary antibodies (Mouse Cardiac Troponin T (Invitrogen) with Rabbit Nkx2.5 (Abcam) or Rabbit CD31/PECAM-1 (Novus Biologicals) at 4 ℃ for 4 days. After washing with PBST, the samples were incubated with secondary antibodies (Goat anti-rabbit Alexa 647, Goat anti-mouse Alexa 594 (Invitrogen)) at 4 ℃ for 2 days. DAPI (4’,6-diamidino-2-phenylindole, Sigma-Aldrich) was added for nuclear staining, and the samples were incubated at 4 ℃ for 24 hours. Finally, the samples were treated with an optical clearing solution and incubated at room temperature for 3 hours. The prepared samples were stored at 4 °C until imaging, which was performed using a Leica TCS SP8 confocal microscope. Images were analyzed using Fiji software.

### Electrophysiological recordings

The Intan RHD recording system was used to record the electrophysiological signals from the samples. Devices were connected to Intan RHD recording headstages, with a platinum electrode immersed in the culture medium serving as both the ground and reference electrode. Signals were recorded for 2 to 5 minutes in a customized Faraday cage with a sampling rate of 10 kHz.

Raw data files were loaded and converted using a Python script provided by Intan Technologies. Noise was reduced by applying either a 60 Hz low-pass filter (*Butterworth*, 4th order) or a 0.3–150 Hz band-pass filter, followed by Savitzky-Golay filter (*window length=101, polynomial order=3*)^51^. Spikes from each channel were identified using the “*scipy.signal.find_peaks*” function from the SciPy library^52^, with the detection threshold set to 3–5 times the standard deviation from the mean. Waveforms were aligned based on absolute maximum dV/dt (maximum dV/dt), and durations of 1.6 seconds were collected for hiPSC-CM homogeneous cultures, while 0.9-second durations were used for hiPSC-CM/hiPSC-EC heterogeneous co-cultures to accommodate their faster beating rates. Further waveform analysis utilized the Python package Scanpy^53^. Waveforms from all channels and recording stages were projected into a 2D representative space using Uniform Manifold Approximation and Projection (UMAP) for dimensional reduction^36^, followed by *Leiden* clustering to assess spike similarities across channels and stages^35^. Representative waveforms from each channel were averaged to generate single-stage waveforms, which were then analyzed in GraphPad Prism for waveform features such as amplitude, dV/dt, and field potential duration (FPD). Waveforms that did not exhibit significant FPD peaks were excluded from the FPD analysis.

Electrophysiological features were extracted using methods adapted from the previous report^24^. For each channel, waveforms of 1.6 seconds (or 0.9 seconds for hiPSC-CM/hiPSC-EC co-cultures) were aligned at the maximum dV/dt and averaged to produce representative waveforms. These representative waveforms were denoised using PyWavelets (*wavelet=‘db6’, level=3, factor=5*)^54^, and 62 feature points were extracted through downsampling. Specifically, 22 feature points were obtained by downsampling the entire waveform (zoomed-out binning), while an additional 40 feature points were extracted from a 0.1-second segment centered around the maximum dV/dt (zoomed-in binning) and concatenated.

### *In situ* sequencing

*In situ* sequencing of RNA in tissue-electronic hybrids were performed following previous methods with modifications^24,25^. The tissue-electronic hybrids were fixed with 4% PFA for 15 minutes then permeabilized with 2 mL -20 °C methanol (Sigma-Aldrich), incubated at -80 °C overnight. The sample was then quenched with PBSTR (1X PBS with 0.1% Tween-20, 0.1 U/µL SUPERase inhibitor (Thermo Fisher)) with 1% yeast tRNA (Invitrogen), 100 mM Glycine (Sigma-Aldrich) for 5 minutes. The sample was then incubated in the 1X hybridization buffer (2X SSC (Sigma-Aldrich), 10% formamide (Calbiochem), 1% Tween-20, 20mM Ribonucleoside Vanadyl Complex (RVC, New England Biolabs), 0.1 mg/mL yeast tRNA, 0.2% SDS (Calbiochem) and the SNAIL oligo pool (Integrated DNA Technologies) with 10 nM per oligo) at 40 °C for 24 hours. The sample was then washed with PBSTR for twice, 20 minutes each at 37 °C then PBSTR with 4X SSC for 20 minutes at 37 °C. Then the samples were washed with PBSTR three times, 5 minutes each. Next, the samples were incubated in the ligation mixture (0.1 Weiss U/µL T4 DNA ligase (Thermo Fisher), 0.2 U/µL SUPERase inhibitor, 0.5 mg/mL Bovine serum albumin (BSA, Invitrogen)) at room temperature overnight, washing with PBSTR for three times, 5 minutes each. The sampled were then incubated in the RCA mixture (0.2 Weiss U/µL Phi29 DNA polymerase (Thermo Fisher), 0.2 U/µL SUPERase inhibitor, 0.5 mg/mL BSA, 20 µM 5-(3-aminoallyl)-dUTP (Invitrogen), 250 µM dNTP mixture (Invitrogen)) for 6 hours then washed with PBST (1X PBS with 0.1% Tween-20) for three times, 5 minutes each. Subsequently, the samples were modified with 25 mM MA-NHS (Methylacrylic acid NHS ester, Sigma-Aldrich) at room temperature for 1 hour, washed with PBST three times and incubating in the monomer solution (4% Acrylamide (Bio-Rad), 0.2% Bis-acrylamide (Bio-Rad), 2X SSC, 0.2% Tetramethylethylenediamine (Sigma-Aldrich)) at room temperature for 15 minutes. The solution was removed and replaced with 30 µL of monomer solution with 0.2% ammonium persulfate, covered with the Gel Slick (Lonza) coated coverslip and polymerized under nitrogen for 1 hour. The samples were then digested with 0.2 mg/mL proteinase K (Invitrogen) at 37 °C overnight then washed with PBST for three times, 10 minutes each. The samples were then treated with 250 U/mL Antarctic phosphatase (New England Biolabs) for 30 minutes before sequencing. For the sequencing reaction, the samples were treated with stripping buffer (60% formamide with 0.1% Triton-X) for twice, 10 minutes each, followed by PBST for three times, 5 minutes each. Then the samples were incubated in the sequencing mixture of different rounds (1:30 T4 DNA ligase, 1:100 BSA, 10 µM reading probes (Integrated DNA Technologies) and 5 µM decoding probes (Integrated DNA Technologies)). DAPI (Sigma-Aldrich) staining was performed according to the manufacturer’s protocol. The imaging was performed on Leica TCS SP8 confocal microscope with 25X water immersion objection with 0.95 N.A.

### Data analysis

#### Imaging data analysis

Image processing and analysis of *in situ* sequencing data were conducted following established protocols. Raw fluorescence images underwent deconvolution using Huygens Essential software. Subsequently, contrast matching was performed using MATLAB’s “*imhistmatchn*” function, followed by background noise suppression via tophat filtering. Global registration of images from five rounds was achieved using 3D fast Fourier transformation to maximize correlation coefficients, compensating for three-dimensional offsets. Further alignment was accomplished through non-rigid registration using MATLAB’s “*imregdemons*” function.

Individual amplicons were identified from first-cycle images using MATLAB’s “*imregionalmax*” function. Gene identity was determined by analyzing the dominant color within a [1, 1, 1] voxel volume surrounding each amplicon’s centroid and referencing the gene codebook. Cell segmentation was performed on the resulting gene amplicon point clouds using ClusterMap^34^.

Single-cell expression analysis utilized the Python package Scanpy. Quality control measures included filtering out cells with fewer than 5 gene counts or 5 unique genes and subsequently integrating with single cell RNA sequencing (scRNA-seq) data we previously published^23,55,56,57^. Data normalization involved log-transformation (log(x+1)) and scaling to unit variance with zero mean. Batch correction across sample conditions was implemented using the “*scanpy.pp.combat*” function prior to downstream analyses^58^.

#### Electrode segmentation and recorded cell identification

Device components, including the graphene/PEDOT:PSS electrode, SU-8 passivation layers, and gold cable, were visualized using reflection mode imaging on a Leica TCS SP8 microscope. The reflection channel data underwent maximum intensity projection (MIP) onto the x-y plane. Electrode segmentation was performed using the Segment Anything Model (SAM, version sam_vit_h_4b8939) by Meta^37^, with manual annotation of foreground and background regions to generate an electrode mask. This mask was then applied to the 3D reflection images to isolate the electrodes, which were subsequently converted to 8-bit (0-255) grayscale images and reconstructed as point clouds. A 3D convex hull was generated from the reconstructed electrode point clouds. To identify recorded cells, the distances between the centers of individual cells and the center of the convex hull were calculated. Cells closest to the electrode side of the convex hull on the electrode side were identified as recorded cells.

#### Niche analysis

For niche analysis, the neighborhood composition for each segmented cell derived from the *in situ* sequencing dataset was computed after cell typing. Based on previous published methods, a radius of 45 microns from the centroid of each cell was used to define the local microenvironment to include ∼2 layers of cells from the center^34^. Within this radius, the composition of neighboring cell types was quantified, resulting in a cell-by-cell-type composition matrix. This matrix was subsequently subjected to UMAP to obtain low-dimensional embeddings that capture the local neighborhood composition of each cell. The resulting embeddings were clustered using the *Leiden* algorithm, enabling the identification of distinct niche types based on shared local neighborhood structures. Two samples were used, both exhibiting a similar distribution of cardiomyocyte states and surrounding cell types (Extended Data Fig. 8). The primary cell niches identified near electrodes differed between samples due to positioning of electrodes within the heterogeneous co-culture and the extent of hiPSC-EC diffusion into the hiPSC-CM-only region. The samples were treated as comparable for further analyses.

#### Pseudotime analysis

Pseudotime analysis was conducted using the *Slingshot* package^43^ to infer the temporal progression of gene expression or electrophysiology in electrically recorded cells. Gene expression profiles or electrophysiological features were projected into 2D UMAP embeddings. Cluster labels derived from *Leiden* clustering were utilized to guide the pseudotime inference. The UMAP embeddings and cluster labels were converted into R-compatible formats and processed through the Slingshot algorithm, which fitted a principal curve to estimate pseudotime trajectories. The resulting pseudotime values were subsequently normalized to a range of [0, 1] for comparability.

## Acknowledgement

We acknowledge Xin Tang for assistance with initial electrophysiological signal processing. J. Liu acknowledges the support from NIH/NIMH 1RF1MH123948; NIH/NIDDK 1DP1DK130673; NSF ECCS-2038603; NIH/NLM 5R01LM014465; and NIH/NHLBI 1R33HL175683. J. Lee acknowledges the support from Kwanjeong Educational Foundation.

## Author contribution

J. Lee and W.W equally contributed to this work. J. Liu, J. Lee, and W. W. conceived the idea and designed the experiments. J. Lee fabricated and characterized the electronics. W. W., Q. L., and R. Liu differentiated and cultured cells. W. W. performed cell/device integration. J. Lee and W. W. conducted electrophysiological recordings, which J. Lee analyzed. J. Lee, W. W., and Z. L. performed immunohistostaining experiments. W. W. and Z. L. conducted the *in situ* sequencing experiments. W. W. analyzed the *in situ* sequencing data with critical input from Z. T. J. Lee and W. W. analyzed the multimodal data. J. A. and R. T. Lee provided the iPSC-derived ECs. J. Lee, W. W., and J. Liu wrote the manuscript. All authors contributed to the writing of the manuscript. J. Liu supervised the study.

## Data availability

The source data supporting the findings of this study are available within the paper.

## Competing interests

J. Liu is cofounder of Axoft Inc.

## Figures and Figure Captions

**Extended Data Fig. 1:**
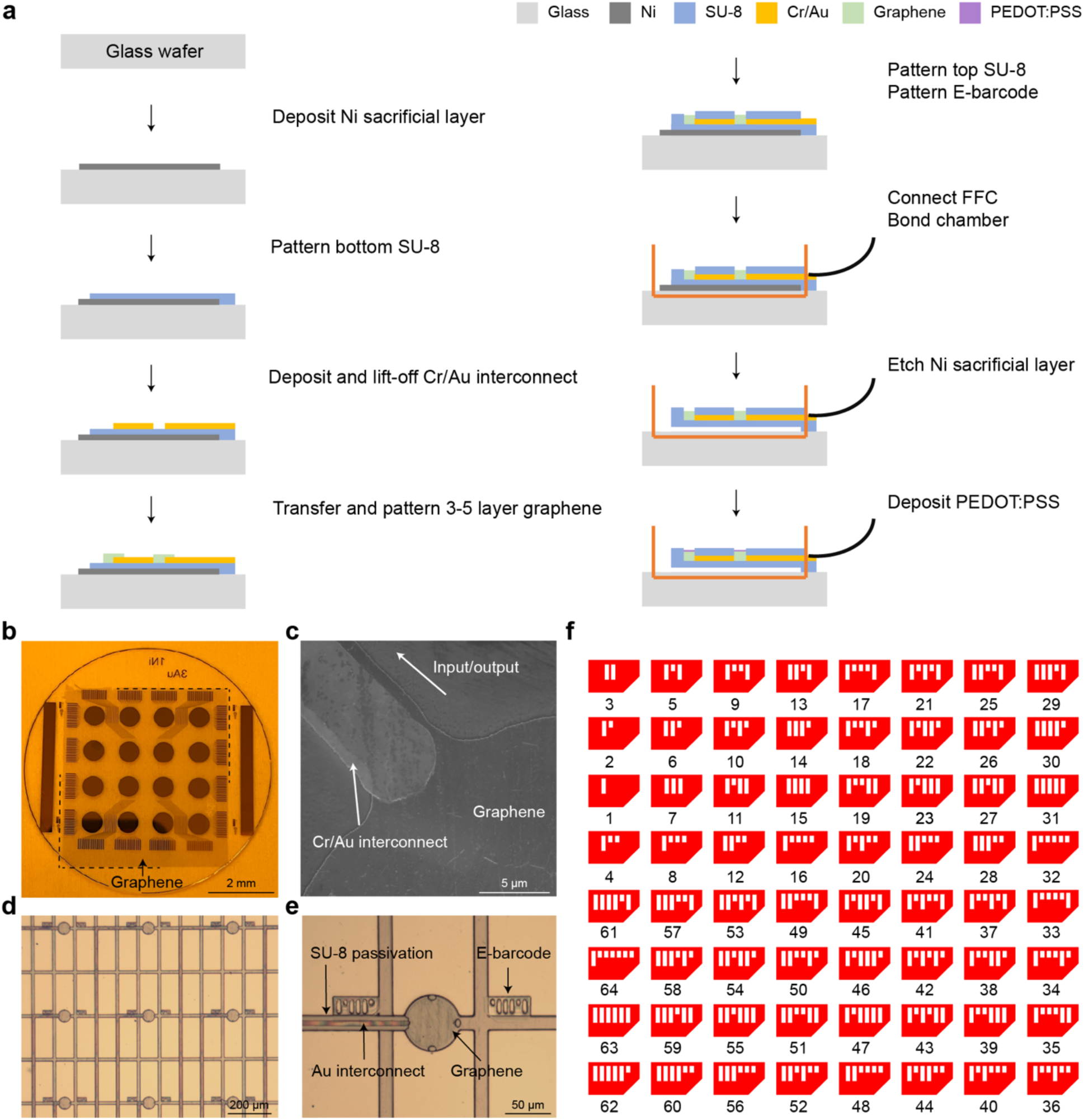
Fabrication of stretchable mesh nanoelectronics with transparent electrodes. **a**, Schematic overview of the fabrication process. **b**, Photograph of graphene transferred onto the devices fabricated on a 3-in glass wafer. **c**, SEM image of graphene on metal interconnect before top SU-8 passivation. Graphene covers part of interconnects, forming an electrically conductive path to I/O. **d**, Optical image of a representative mesh electronics array. **e**, Optical image of a representative graphene electrode with a lithographically patterned fluorescent E-barcodes. **f**, Design layout of E-barcodes for labeling electrodes in the array.

**Extended Data Fig. 2:**
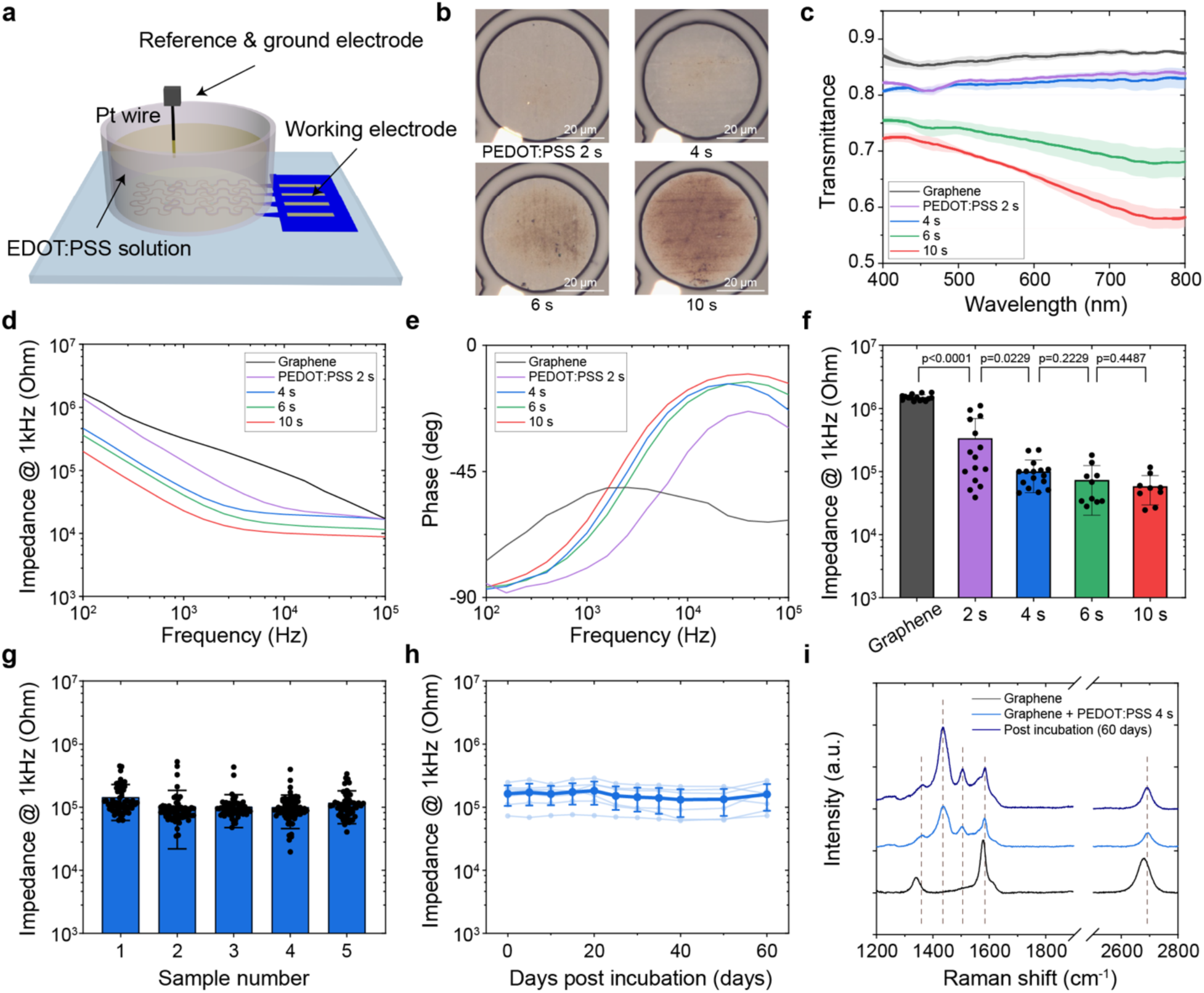
Electrochemical deposition of PEDOT:PSS on graphene electrodes. **a**, Schematic diagram of the setup used for PEDOT:PSS electrochemical deposition. **b**, Optical images of representative graphene electrodes deposited with PEDOT:PSS at varying deposition time. **c**, Transmittance spectra of representative graphene/PEDOT:PSS electrodes from 400 to 800 nm. Data are presented as mean ± S.D., n=5. **d**–**e**, Amplitude (**d**) and phase angle (**e**) of electrochemical impedance spectra measured from representative graphene/PEDOT:PSS electrodes across frequencies ranging from 0.1 to 100 kHz. **f**, Electrochemical impedance at 1 kHz of graphene/PEDOT:PSS electrodes with varying deposition time. Data are presented as mean ± S.D., p-values were calculated from Welch’s two-tailed unpaired t-test, n=15, 15, 16, 10, 10. Each dot represents the impedance of a single electrode within the sample. **g**, Electrochemical impedance at 1 kHz across electrodes from five representative devices. Data are shown as mean ± S.D., each dot indicates the impedance of a single electrode within the sample. **h**, Impedance stability at 1 kHz over time for seven representative electrodes incubated in 37 °C 1X PBS. Values are presented as mean ± S.D. **i**, Raman spectra of representative graphene and graphene/PEDOT:PSS electrodes before and after 60 days of incubation in 37 °C 1X PBS.

**Extended Data Fig. 3:**
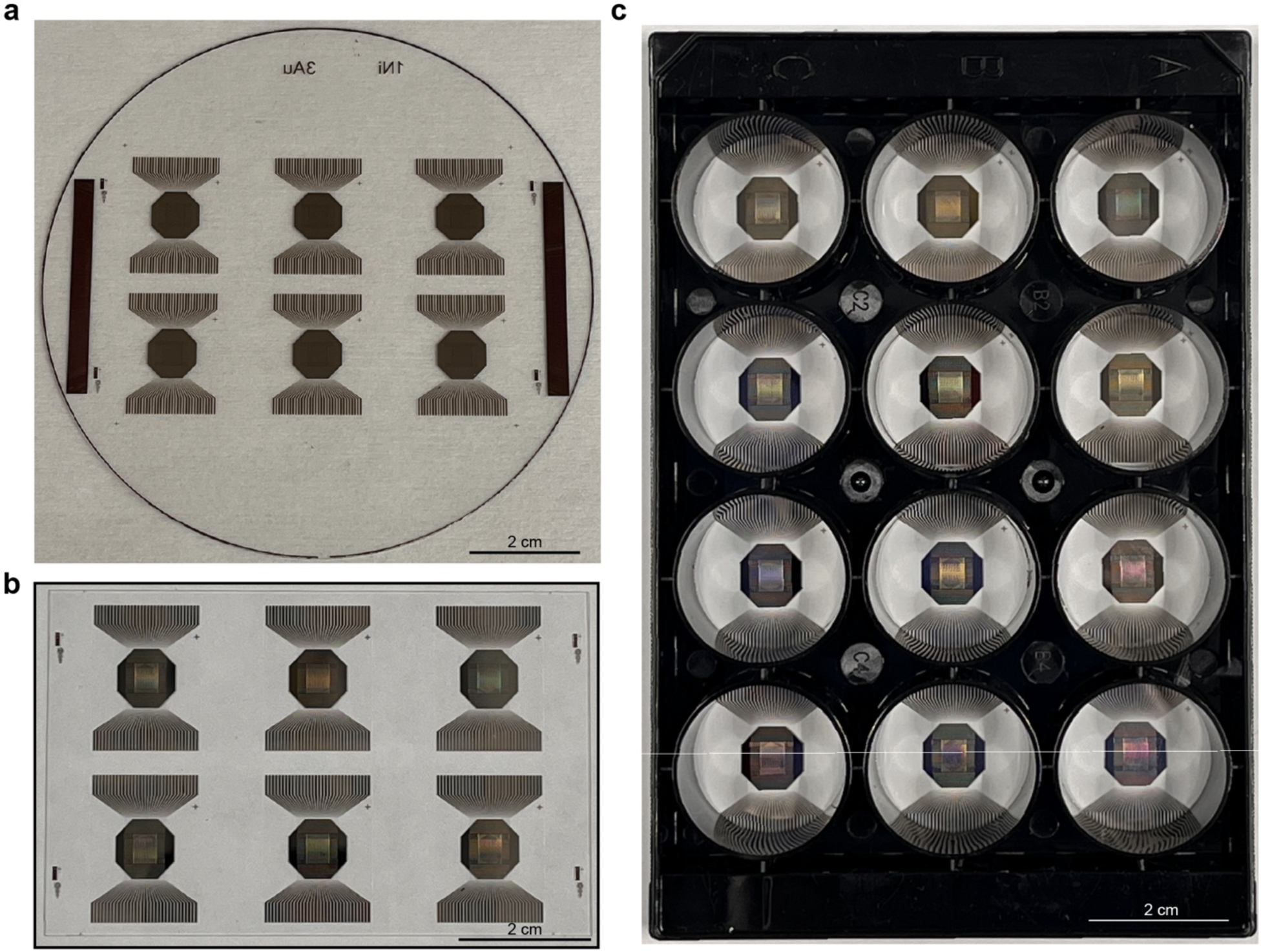
Multiwell plate compatibility for high-throughput *in situ* graphene-seq. **a**, Photograph of devices fabricated on a 4-inch glass wafer, each containing a 64 electrode arrays. **b**, Photograph of devices after post-fabrication processing. **c**, Photograph of devices attached to a 12-well cell plate for tissue culture.

**Extended Data Fig. 4:**
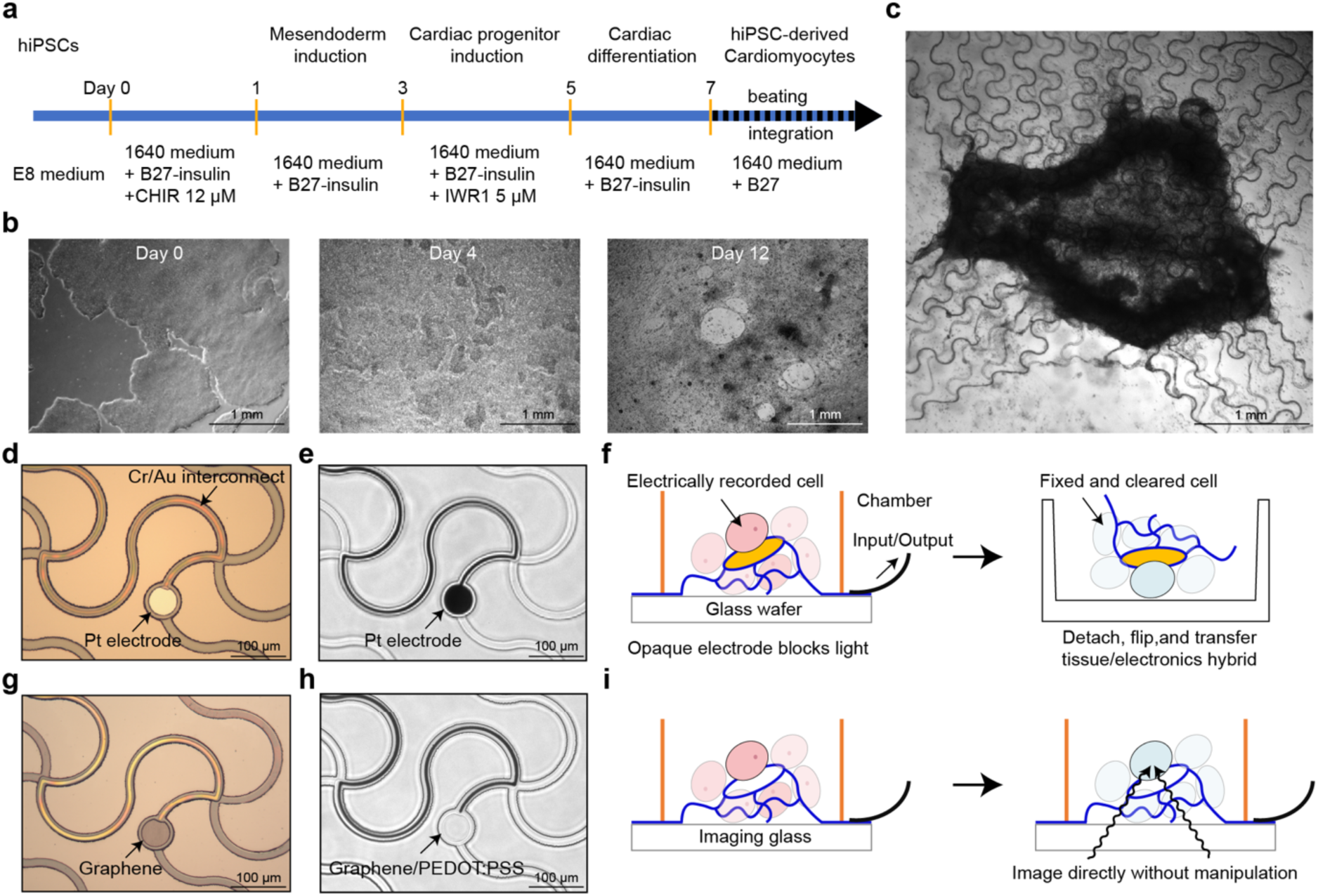
Differentiation and integration of hiPSC-CMs with stretchable and transparent mesh electronics. **a**, Schematic illustrating the protocol for differentiating CMs from hiPSCs. **b**, BF images showing cells at different stages of the differentiation process. **c**, BF image of a 3D cardiac organoids integrated with mesh electronics. **d**–**e**, Optical images of conventional mesh electronics with metal electrodes before (**d**) and after (**e**) being released from the substrate. **f**, Schematic illustrating the additional experimental steps required to image cells on top of opaque metal electrodes. **g**–**h**, Optical images showing graphene electrodes before (**g**) and after (**h**) being released from the substrate and deposited with PEDOT:PSS. **i**, Schematic illustrating how transparent electrodes enable direct imaging of electrically recorded cells.

**Extended Data Fig. 5:**
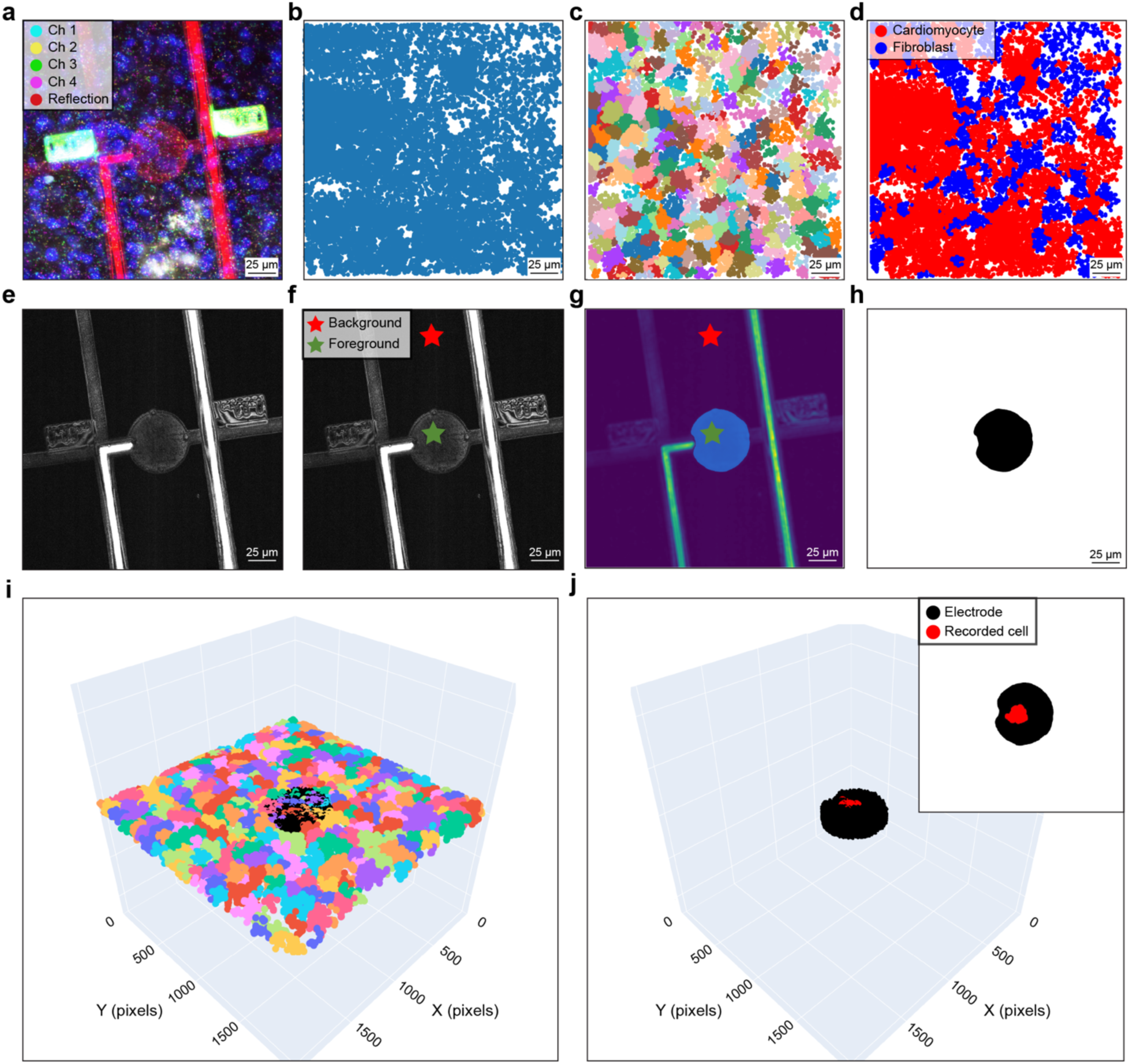
Image processing pipeline for *in situ* sequencing data. **a**–**d**, *In situ* sequencing data processing pipeline. **a**, Representative raw cDNA amplicons (channels 1–4) showing fluorescence barcodes (green) and the device in reflection mode (red). **b**, Registered cDNA amplicons. **c**, Segmented cells visualized using ClusterMap for cell mapping. **d**, Assigned cell types mapped to segmented cells. **e**–**h**, Graphene electrode segmentation pipeline. **e**, Maximum intensity projection of the device in reflection mode. **f**, Segmentation of background and foreground regions to identify electrodes. **g**, Generated electrode mask. **h**, Binary mask of segmented electrodes for downstream analyses. **i**–**j**, Identification of electrically recorded cells. **i**, 3D reconstruction of electrodes with cells color-coded by segmentation. **j**, Identification of electrically recorded cells positioned over the electrode.

**Extended Data Fig. 6:**
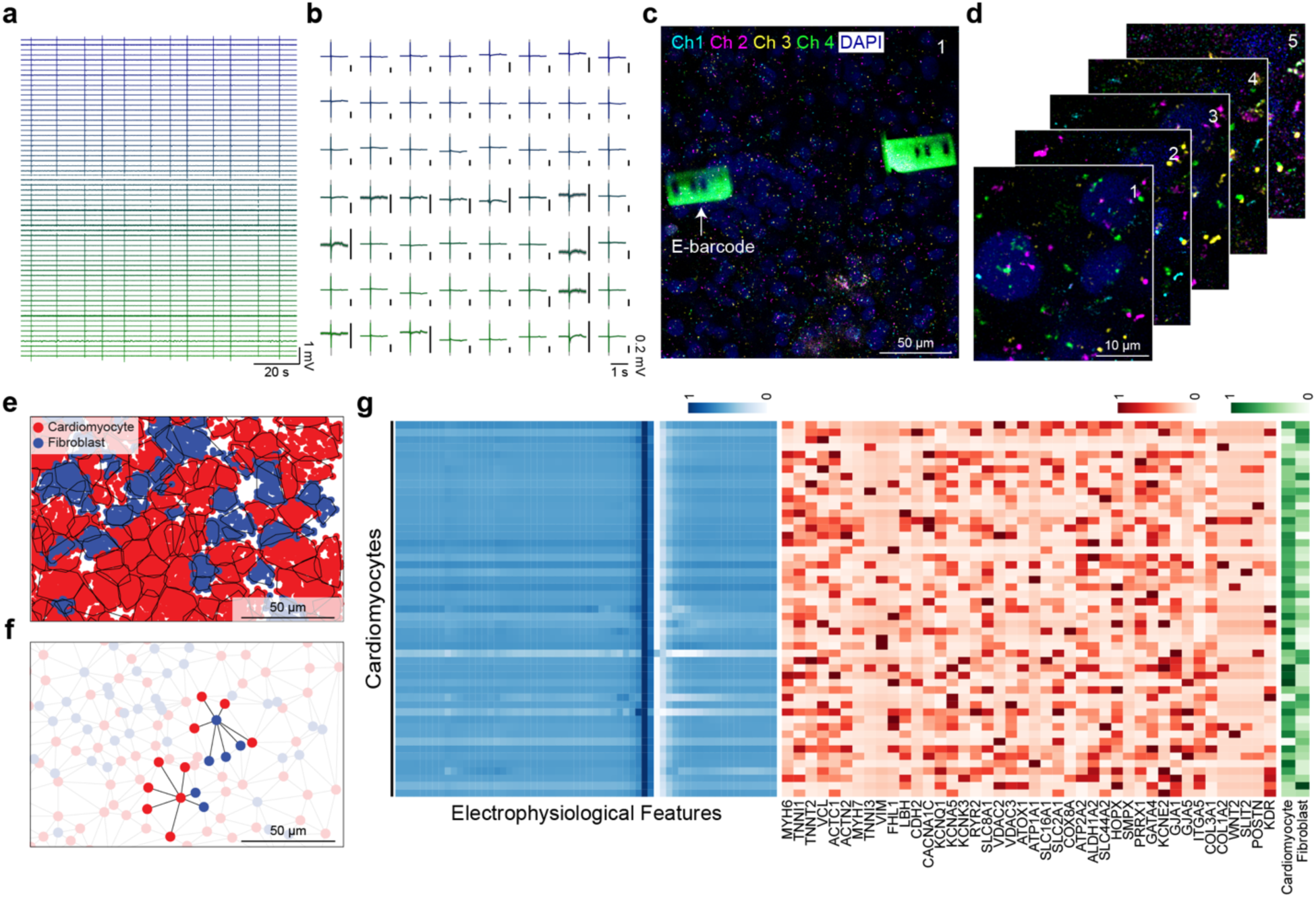
*In situ* graphene-seq of hiPSCs-derived cardiac microtissue. **a**, Representative voltage traces recorded from 64 representative electrodes. **b**, Extracted spike waveforms (mean ± S.D.) corresponding to the recorded signals in (**a**). **c**, Representative image of *in situ* sequencing showing fluorescence channels and E-barcodes. **d**, Zoomed-in view of five cycles of *in situ* sequencing. **e**, Cell-type map showing the spatial distribution of different types of cells within the microtissue. **f**, Cell adjacency networks of a representative cardiomyocyte and fibroblast, illustrating spatial proximity and cellular interactions. **g**, Concatenated multimodal heatmaps of electrophysiological features, gene expression profiles, and local cell density of electrically recorded cardiomyocytes across the microtissue.

**Extended Data Fig. 7:**
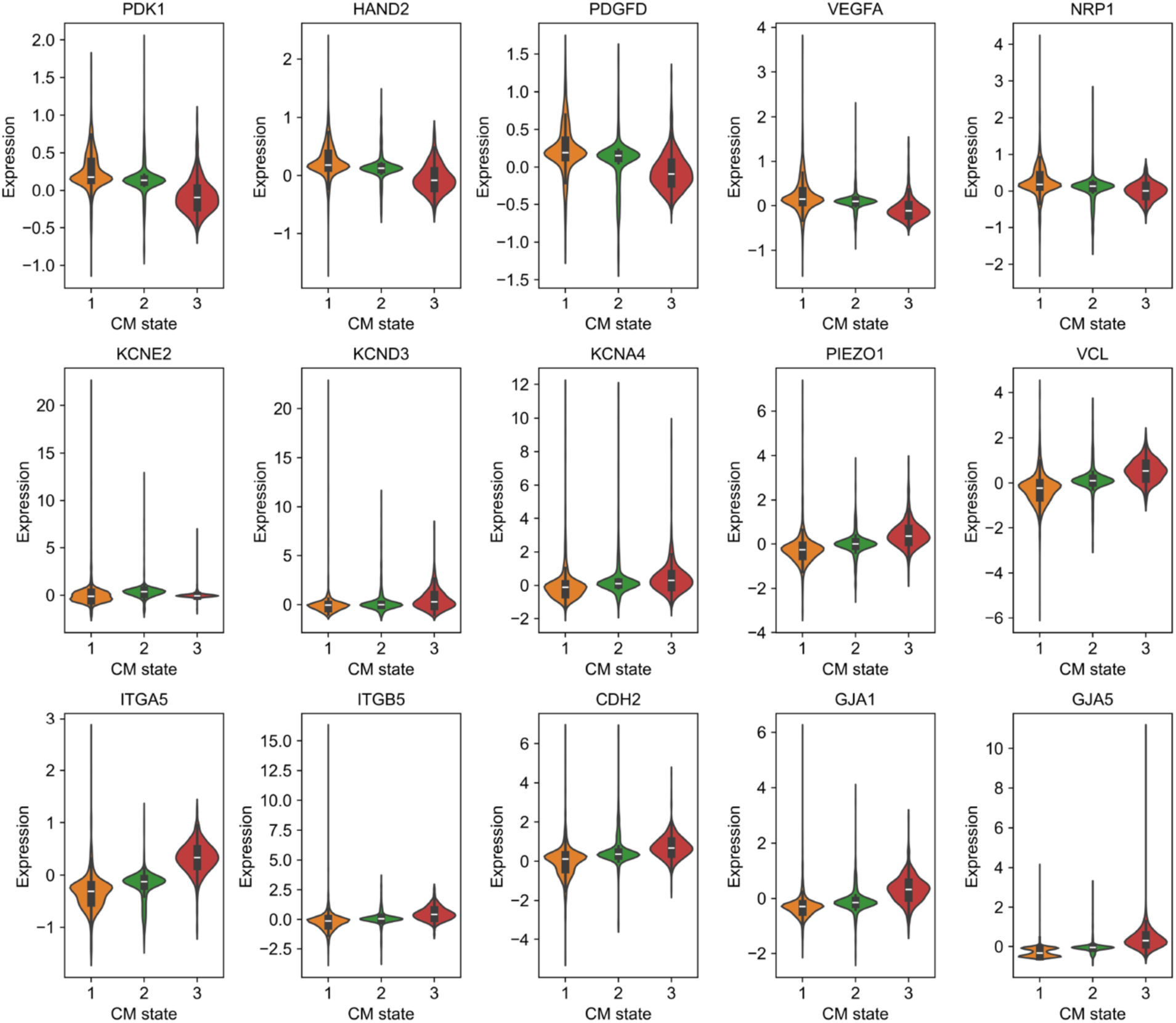
Gene expression profiles of different hiPSC-CM states. Violin plots illustrating the expression levels of selected genes across different cardiomyocyte states. Values are normalized, log-transformed (log(x+1)), and scaled to unit variance and zero mean.

**Extended Data Fig. 8:**
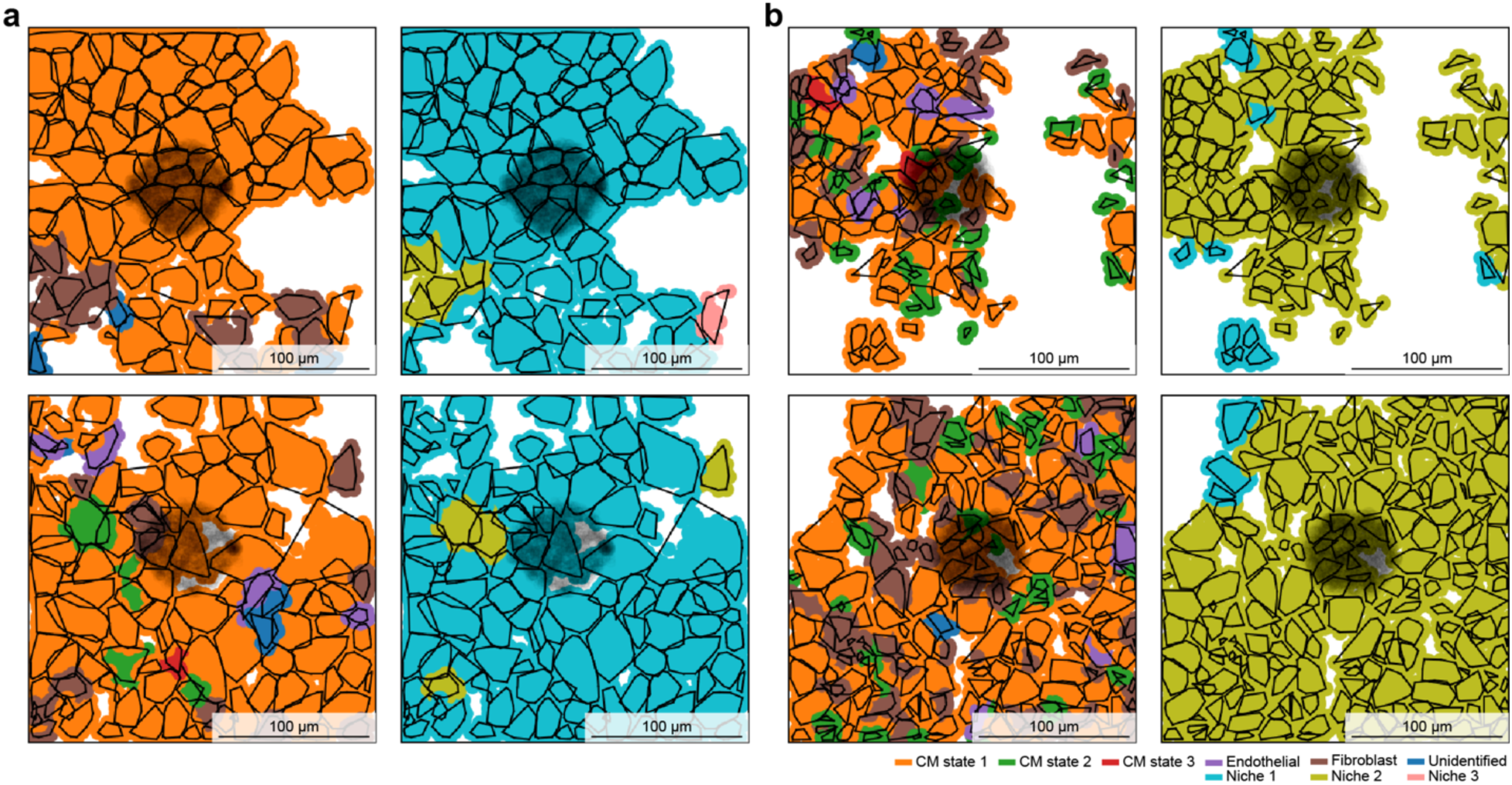
Sample information. **a**–**b**, Representative cell-type maps (left) and niche-type maps (right) of the CM-only regions from hiPSC-CM/hiPSC-EC co-cultured samples. **a**, Sample 1. **b**, Sample 2.

